# Intrinsic temporal tuning of neurons in the optic tectum is shaped by multisensory experience

**DOI:** 10.1101/540898

**Authors:** Silas E. Busch, Arseny S. Khakhalin

## Abstract

For a biological neural network to be functional, its neurons need to be connected with synapses of appropriate strength, and each neuron needs to appropriately respond to its synaptic inputs. This second aspect of network tuning is maintained by intrinsic plasticity; yet it is often considered secondary to changes in connectivity, and mostly limited to adjustments of overall excitability of each neuron. Here we argue that even non-oscillatory neurons can be tuned to inputs of different temporal dynamics, and that they can routinely adjust this tuning to match the statistics of their synaptic activation. Using the dynamic clamp technique, we show that in the tectum of Xenopus tadpoles, neurons become selective for faster inputs when animals are exposed to fast visual stimuli, but remain responsive to longer inputs in animals exposed to slower, looming or multisensory stimulation. We also report a homeostatic co-tuning between synaptic and intrinsic temporal properties of individual tectal cells. These results expand our understanding of intrinsic plasticity in the brain, and suggest that there may exist an additional dimension of network tuning that has been so far overlooked.

## Introduction

It is often assumed, in fields as diverse as connectomics and machine learning, that the main difference between functional and dysfunctional neural networks lies in their connectivity (Takemura, 2014; Hildebrand et al., 2017; Bassett and Sporns, 2017; Reimann et al., 2017). However, biological neurons also demonstrate complex and multifaceted intrinsic tuning (O’Leary et al., 2013; Evans et al., 2015), where cells within a single network having different activation thresholds (Kole and Stuart, 2012), bursting behaviors (Popovic et al., 2011), and inactivation profiles (Bianchi et al., 2012). Dysregulation of intrinsic plasticity affects the network dynamics (Tien and Kerschensteiner, 2018), and can lead to a loss of function (Marcelin et al., 2009). And yet, with the possible exception of oscillatory networks (Marder and Taylor, 2011; Picton et al., 2018), the exact ways in which intrinsic plasticity contributes to network tuning, remain unclear. We know that neurons adjust their spikiness to match the levels of synaptic activation they experience (Aizenman et al., 2003; Titley et al., 2017), but we also know that intrinsic properties can affect more subtle neuronal tuning to different temporal patterns of activation (Azouz and Gray, 2000; Branco et al., 2010; Fontaine et al., 2014; Jarvis et al., 2018; Ohtsuki and Hansel, 2018; Zbili et al., 2019). This begs the question: do neurons use this type of tuning in practice, dynamically adjusting it to the temporal dynamics of their inputs?

In this paper, we investigate whether exposure to sensory stimuli with different temporal dynamics would change intrinsic temporal tuning of sensory neurons in the optic tectum of *Xenopus* tadpoles. The tadpole tectum is an ideal model for exploring this question: it is a highly malleable distributed network of about 10^4^ neurons (Pratt and Khakhalin, 2013), involved in stimulus discrimination and sensorimotor transformations (Dong et al., 2009; Khakhalin et al., 2014). In development, tectal neurons acquire diverse phenotypes, that are then shaped by sensory experiences (Xu et al., 2011; Ciarleglio et al., 2015). Circuits in the tectum can learn and reproduce temporal patterns to which they were exposed (Pratt et al., 2008): a property that could in principle be achieved through synaptic changes alone (Lukoševičius and Jaeger, 2009), but which may be easier to achieve through intrinsic temporal tuning (Narayanan and Johnston, 2008; Beatty et al., 2014). Finally, tectal neurons exhibit strong Na channel inactivation, which seems to play a role in collision detection (Jang et al., 2016), can support temporal tuning (Clay et al., 2012; Fontaine et al., 2014; Zbili et al., 2019), and is a known target for plasticity mechanisms (Bianchi et al., 2012).

Specifically, we ask three questions about the properties of intrinsic plasticity in tectal networks. First, we test whether the intrinsic temporal tuning to either faster (more synchronous) or slower (asynchronous) inputs would change in response to sensory stimulation. Then, we ask whether intrinsic temporal tuning of individual tectal cells is coordinated with the typical length of synaptic currents they receive. Finally, we try to identify the mechanisms that may underlie temporal tuning variability.

In our previously published large-scale census of tectal cells (Ciarleglio et al., 2015), we observed no signs of intrinsic temporal tuning (figures 2, 4), and no interaction between intrinsic and synaptic phenotypes. We argue however, that the current clamp protocols used in earlier studies were not adequate for the task, so in this paper we rely on a dynamic clamp technique. The main benefit of the dynamic clamp, compared to commonly used current step injections, is that it allows a more realistic simulation of neuronal responses to synaptic conductances (Prinz et al., 2004a). In dynamic clamp, the current injected into the cell is adjusted in real-time, based on a predefined formula that depends on time and cell membrane potential. This means that with dynamic clamp recordings we can excite a neuron in a controlled manner, but still allow both its voltage and transmembrane currents to change, preserving feedback interactions between active currents and transient voltage-gated channel inactivation (Ma and Koester, 1996; Zbili et al., 2019), which is an important mechanism for temporal tuning (Branco et al., 2010; Platkiewicz and Brette, 2011). As a second methodological innovation, instead of relying on one type of sensory stimulation (Ciarleglio et al., 2015), here we used five different stimulation protocols, and compared their individual effects on intrinsic tuning.

We show that the intrinsic plasticity of tectal neurons supports temporal selectivity, which is shaped by sensory experience, and is coordinated with the typical duration of synaptic inputs received by each cell. Moreover, the use of different sensory modalities for stimulation gave us an insight into an unrelated, but intriguing question of multisensory integration in the brain (Deeg et al., 2009; Felch et al., 2016; Truszkowski et al., 2017), as for the first time we were able to look at tectal retuning in response to multisensory experience in freely behaving tadpoles.

## Results

All analysis scripts and summary data for every cell can be found at: https://github.com/khakhalin/Dynamic-clamp-2018

### Changes in excitability in response to sensory stimulation

First we checked whether our stimulation protocols caused any changes in intrinsic excitability of tectal neurons. From previous studies, we knew that in tadpoles exposed to LED flashes, tectal neurons became more excitable (Aizenman et al., 2003; Ciarleglio et al., 2015). However, the stimuli we used in the present study were weaker, and similar to those used in behavioral experiments (Khakhalin et al., 2014; James et al., 2015; Truszkowski et al., 2017). We presented a checkerboard pattern that inverted once a second, for four hours; either instantaneously (dubbed “Flash”; Figure 1C left), or with a slow transition over the course of a second (old black squares shrank to white, while new black squares grew from old white squares, dubbed “Looming”; Figure 1C right).

**Figure 1.**
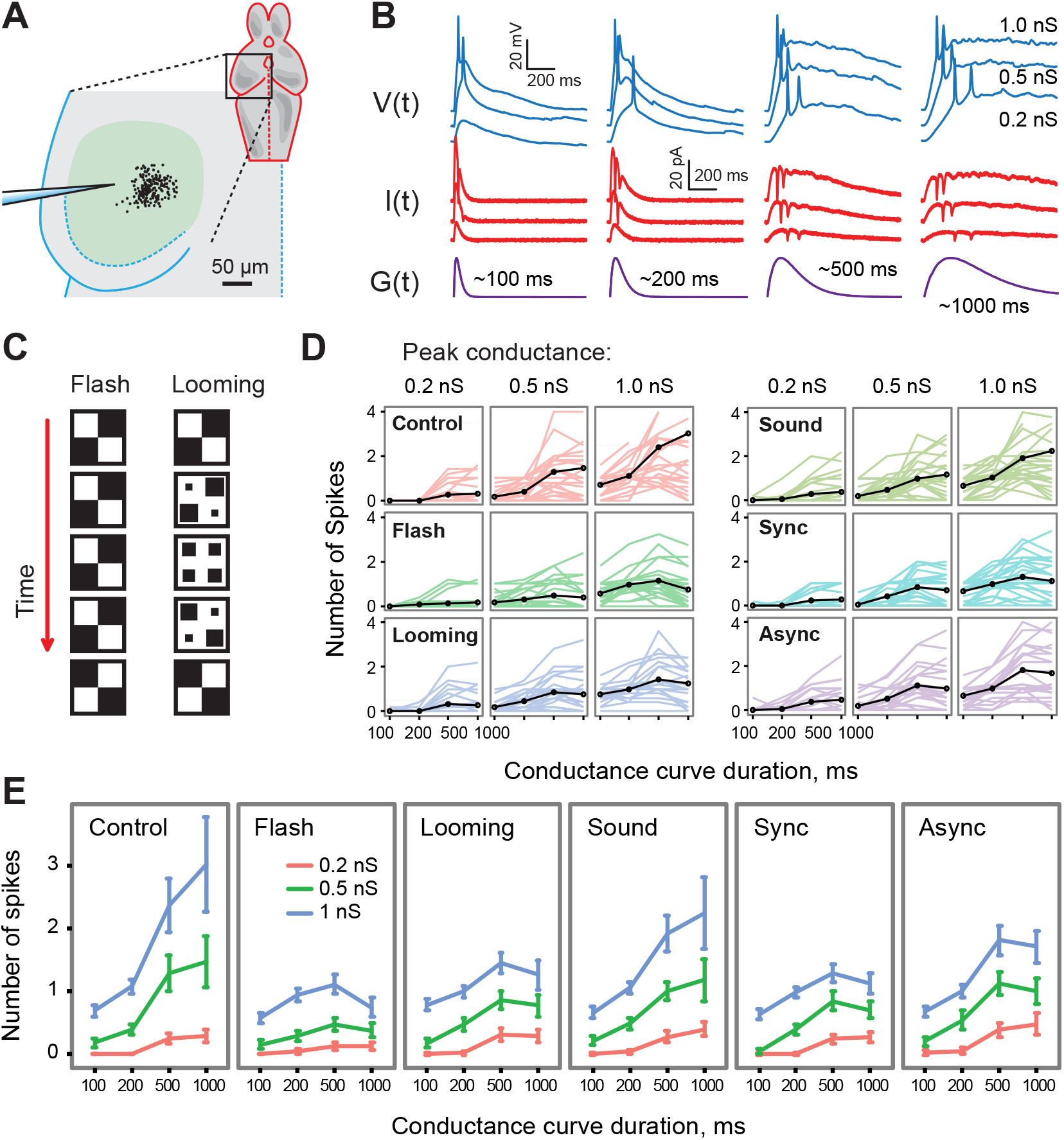
Overview of experimental design and summary of dynamic clamp results. (**A**). Positions of tectal neurons that were recorded. (**B**). Sample data from a dynamic clamp experiment. Bottom row: the dynamics of conductances G(t) of four different durations simulated by the dynamic clamp system. Middle row: the currents I(t) dynamically injected into a cell based on conductances of 4 different durations and 3 different amplitudes. Top row: resulting voltage traces V(t) that were recorded and analyzed. (**C**). A schematic of visual conditioning in “Flash” (left) and “Looming” (right) groups. (**D**). The number of spikes produced by all neurons in all experiments, split by input peak conductance, and plotted against conductance duration. Black lines show respective averages. (**E**). A summary of data from D, presented as averages and 95% confidence intervals.

After conditioning, we excised the brain, performed whole cell patch clamp on tectal neurons (Figure 1A), and counted spikes produced in response to dynamically simulated synaptic conductances (Figure 1B). We used conductances of 4 different durations (100, 200, 500, and 1000 ms), and 3 different amplitudes (peak conductances of 0.2, 0.5, and 1.0 nS), matching the range of synaptic currents observed in tectal circuits *in vivo* (Xu et al. 2011; Khakhalin et al. 2014; Ciarleglio et al. 2015; see Methods). Based on prior research (Aizenman et al., 2003; Ciarleglio et al., 2015), we expected neurons exposed to instantaneous checkerboard inversions (flashes) to become more excitable, but surprisingly, they were on average less spiky, and generated 0.4±0.4 spikes, across all types of dynamic clamp inputs, compared to 0.9±1.0 spikes in control (Figure 1D, left column; F(1,677)=30.4, p=5e-8, n cells=28, 29). (Here and below, F-values are reported for a multivariate fixed effects analysis of variance with selected interactions, where cell ID is used as a repeated measures factor; see Methods for a detailed description). Neurons exposed to looming transitions also spiked less than control neurons (0.6±0.4 spikes; F(1,629)=9.6; p=0.002; n=28, 25), but more than those exposed to instantaneous “flashes” (F(1,641)=3.7; p=4e-6; n=29, 25). This was likewise unexpected, as compared to flashes, looming stimuli are known to elicit stronger tectal responses (Khakhalin et al., 2014; Khakhalin, 2019), yet their effect on neuronal excitability was weaker, seemingly in contradiction with the principle of homeostatic plasticity (Pratt and Aizenman, 2007; Turrigiano, 2007).

We then mapped the amplitude tuning of neurons (their amplitude transfer function, or gain), by looking at how an increase in transmembrane conductance translated into increased spike output. Compared to control, neurons from animals exposed to visual stimuli had a flatter amplitude tuning curve (less of a difference between columns of Figure 1D, or between curves in each individual panel of Figure 1E), as they did not increase their spiking as fast in response to larger conductances (F(1,677)=15.9, p=8e-5; and F(1,629)=8.3, p=0.004 for flash- and looming respectively, compared to control). The flattening of response curves seemed slightly more pronounced for flashes than for looming (F(1,641)=3.5; p=0.06).

These results show that prolonged stimulation had an effect on intrinsic excitability, but its direction was opposite to what was previously described (Aizenman et al., 2003; Ciarleglio et al., 2015), as neurons became less excitable. Moreover, while looming stimuli are known to be more salient, both behaviorally and physiologically (Khakhalin et al., 2014), they had a weaker effect on neuronal excitability in comparison to less salient flashes. We conjecture that the difference in the direction of change is due to our stimuli being weaker than those used in earlier studies (see Discussion), and we further explore the difference between flashes and looming stimuli below.

### Changes in intrinsic temporal tuning

We then examined whether different types of sensory activation would differently reshape intrinsic temporal tuning in tectal neurons. We knew from earlier studies that fast tectal responses can be evoked with instantaneous flashes, while looming stimuli cause slower, and more prolonged tectal activation (Khakhalin et al., 2014; Khakhalin, 2019). As overall changes in intrinsic properties seemed homeostatic (increased activation led to reduced spiking), by the same logic, one could expect that exposure to flashes would selectively suppress spiking in response to fast synaptic inputs. On the other hand, one could argue that exposure to shorter stimuli could make neurons better adjusted to working with fast synaptic currents (Stemmler and Koch, 1999), similar to changes described for synaptic plasticity (Aizenman and Cline, 2007) and recurrent activity in the tectum (Pratt and Aizenman, 2007; Shen et al., 2011).

We found that compared to control, neurons exposed to flashes and looming stimuli had different response curves when tested across input conductances of different lengths (F(1, 677)=25.1; p=7e-7; and F(1,629)=12.0; p=6e-4 respectively). In control neurons, longer inputs (500 and 1000 ms) typically evoked stronger spiking than shorter inputs (100 and 200 ms), but neurons from stimulated animals had flatter tuning curves, with a plateau, or even a decrease in spike output for longer conductance injections (Figure 1E). In other words, while control neurons preferred longer inputs (responded to them with more spikes), stimulated neurons developed a preference for shorter synaptic inputs, and this change was more pronounced in neurons exposed to flashes than in those exposed to looming stimuli (F(1,641)=7.5; p=0.006). This suggests that the change in overall intrinsic excitability, and the change in temporal tuning, followed two different kinds of logic. The overall excitability was homeostatic, as neurons became less excitable in response to stronger stimulation. The temporal retuning however can be better described as “adaptive”, as neurons exposed to shorter stimuli (flashes) became relatively *more* responsive to shorter stimuli, and less responsive to longer stimuli, making them equipped to process faster, synchronous patterns of activation (Stemmler and Koch, 1999; Fontaine et al., 2014).

### Effects of acoustic and multisensory stimulation

While the optic tectum (homologous to superior colliculus in mammals) is often described as a visual area, it is also involved in heavy multisensory computations (Stein et al., 2014). In tadpoles, it integrates visual information with inputs from mechanosensory, auditory, and lateral line modalities (Deeg et al., 2009; Pratt and Aizenman, 2009; Hiramoto and Cline, 2009; Felch et al., 2016; Truszkowski et al., 2017), but the logic of this integration is unclear. We wondered whether acoustic stimuli would reshape intrinsic properties of tectal neurons, and whether this retuning would be similar to that produced by visual stimuli.

To test this question, we exposed tadpoles to four hours of behaviorally salient, startle-inducing “click” sounds (James et al., 2015; Truszkowski et al., 2017), provided at the same frequency (every second) as visual stimuli in the first set of experiments. We found (Fig 1D, E) that exposure to these sounds (group “Sound”) did not lead to significant changes in either average number of spikes (0.8±0.7 spikes; F(1,689)=1.7; p=0.2; n=28, 30), amplitude transfer function (F(1,689)=2.0; p=0.2), or temporal tuning curve (F(1,689)=2.0; p=0.2). This may suggest that acoustic stimuli did not activate tectal circuits strongly enough during conditioning, despite being more behaviorally salient (at the onset of stimulation, acoustic clicks evoked startle responses in about 50-80% of cases, compared to 5-10% for checkerboard inversions (James et al., 2015; Truszkowski et al., 2017)). This was not necessarily surprising, as mechanosensory and visual inputs have different cellular targets in the tectum (Pratt and Aizenman, 2009; Felch et al., 2016; Truszkowski et al., 2017), potentially leading to different recruitment of tectal excitatory and inhibitory circuits. Acoustic stimuli may also be inherently weaker than visual stimuli in triggering plasticity effects in tectal neurons, as they arrive at different compartments within the dendritic tree (Hiramoto and Cline, 2009; Deeg et al., 2009), which may define their influence on neuronal plasticity (Richards and Lillicrap, 2019).

We then combined visual and acoustic stimuli in two different ways and looked at the effects of multisensory stimulation on the intrinsic properties of tectal neurons. For some animals, we synchronized the instantaneous checkerboard inversions (flashes) with sound clicks (dubbed “Sync”), while for others, we staggered visual and acoustic stimuli by half a period (500 ms; dubbed “Async”). We found (Fig 1D, E) that multisensory stimulation did not suppress excitability of tectal neurons as strongly as visual stimulation alone (0.6±0.4 spikes, F(1,689)=11.2, p=8e-4; and 0.7±0.6 spikes, F(1,665)=41.7, p=2e-10, for Sync and Async respectively, compared to Flash, across all testing conditions). Moreover, both temporal and amplitude tuning curves in multisensory groups were less flat than in the Flash group, which was especially noticeable for the Async group (for amplitude tuning: F(1,689)=1.3, p=0.3, and F(1,665)=2.4, p=0.02 in Sync and Async groups respectively; for temporal tuning: F(1,689)=9.3, p=0.002, and F(1,665)=22.8, p=2e-6 respectively). This suggests that while on their own sound clicks had little effect on tectal excitability, when added to visual flashes, they negated some of the retuning effects that visual stimulation would have had.

### Changes in average neuronal tuning, and tuning variability

To better visualize and interpret differences in neuronal tuning after different types of stimulation, we quantified three aspects of intrinsic excitability (average spikiness, amplitude tuning, and temporal tuning) with one value per neuron (see Methods). We used the mean number of spikes across all conditions as the measure of “spikiness”; the linear slope of the number of spikes as a function of conductance amplitude as the measure of “amplitude tuning”, and the quadratic term of the curvilinear regression for the number of spikes as a function of input duration as the value to characterize the “temporal tuning” of each neuron (Figure 2A). These “tuning coefficients” capture the character of tuning curves for each neuron (Figure 2A). All three parameters differed across experimental groups: F(5,160)=3.1, p=0.01 for average spikiness (Figure 2C; see Methods for model description); F(5,160)=4.8, p=4e-4 for amplitude tuning (Figure 2B); and F(5,160)=3.6, p=4e-3 for temporal tuning (Figure 2C).

**Figure 2.**
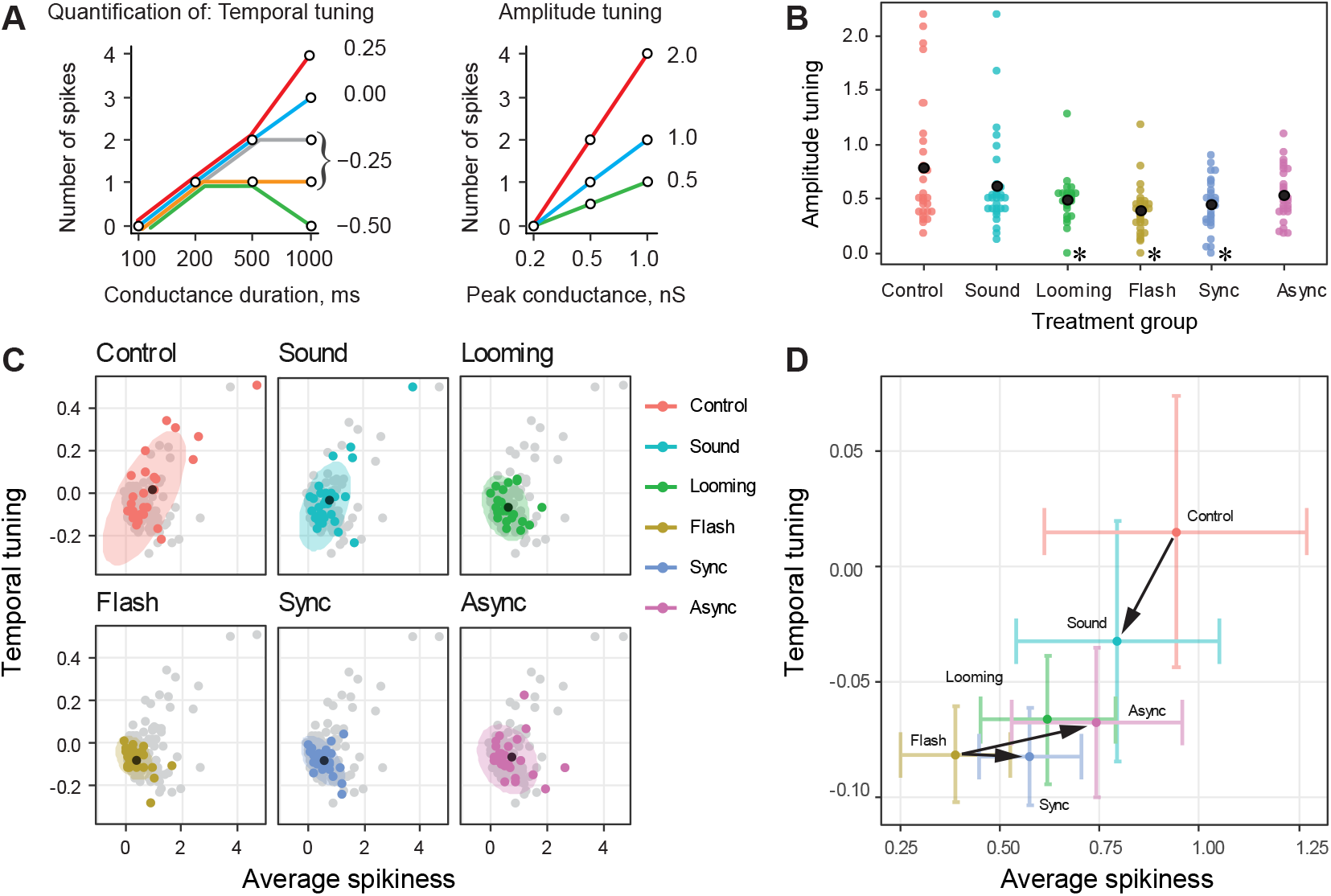
Quantification of changes in temporal tuning in response to sensory experience. (**A**). An illustration of how “Temporal tuning” and “Amplitude tuning” values were calculated. For the temporal tuning measure, the value of zero corresponds to linear dependency (blue line), positive values - to an accelerating, supralinear curve (red), and negative values - to plateau- and hill-shaped curves (gray, orange, green). For amplitude tuning, higher values correspond to faster increase in spiking with increased conductance. (**B**). Amplitude tuning of neurons across different experimental groups (stars show t-test p<0.05 compared to control). (**C**). Temporal tuning and spikiness of neurons in different experimental groups. Neurons from each target group are shown in color, while all neurons from all groups are shown in light gray, as a reference; groups means are shown as black dots; ellipses represent 95% normal confidence regions. Two outliers (top right corner) are brought within the axes limits. (**D**). Same data as in (C), shown as averages for each group, with 95% confidence intervals. Black arrows show the effects of sound clicks, when they were added to control (red), and when they were added to “Flashes” (brown), to form two types of multisensory stimuli.

A visual comparison of neuronal tuning in different experimental groups (Fig 2C) shows that acoustic stimulation had opposite effects when provided on its own (without visual stimulation), compared to when it was added to visual flashes. Compared to control neurons, cells exposed to sound clicks seemed to have lower amplitude (Cohen’s d= −0.29) and temporal (d= −0.27) tuning coefficients (Hotelling t-squared test p=0.2). When sounds were added to flashes, however, both tuning coefficients increased (d=0.65 and 0.26 for amplitude and temporal tuning respectively, “Async” compared to “Flash”), making neurons more similar to control neurons (Hotelling test p=0.04). Thus, acoustic stimulation tended to tune the network in the same direction as visual stimulation when delivered alone (Figure 2D), but it negated the effects of visual stimulation when the two were combined. This effect is likely to be either a consequence of strong inhibitory recruitment during multisensory integration in the tectum (Liu et al., 2016; Hamodi et al., 2016), or a sign of highly non-linear interactions between acoustic and visual inputs within the dendritic trees of individual tectal neurons (Deeg and Aizenman, 2011; Felch et al., 2016; Truszkowski et al., 2017).

Somewhat similarly, looming stimuli seemed to have weaker effects on intrinsic properties, compared to flashes, both in terms of amplitude (d=0.44) and temporal tuning (d=0.22), despite them eliciting stronger responses in-vivo (Khakhalin et al., 2014; Khakhalin, 2019). This suggests that intrinsic retuning depends on stimuli synchronicity, rather than simply the total number of spikes generated by the network, as more synchronous stimuli (Flash, Sync) had stronger effects on tuning than comparable asynchronous stimuli (Async, Looming).

Describing neuronal tuning with only a few variables also allowed us to compare cell-to-cell variability of tuning in different experimental groups. We found that, as it can be guessed from Figure 2D, this variability decreased as neurons were modulated away from the baseline (Bartlett test p=2e-9 for amplitude tuning, p=1e-9 for temporal tuning). Groups that were significantly different in average values also had different variances (F-test with p<0.05), such as Control vs. Flash and Control vs. Looming for both amplitude and temporal tuning. This expands on the findings of our previous study (Ciarleglio et al., 2015), where we showed that prolonged patterned stimulation reduces diversity of tuning profiles in the network, reshaping them according to the spatiotemporal characteristics of the stimulus. We now show that stimuli that reshape the network stronger, also have a more restrictive effect on tuning diversity.

### Changes in synaptic properties

To see whether prolonged sensory stimulation affected synaptic inputs received by tectal neurons, we recorded evoked excitatory postsynaptic currents in response to optic chiasm stimulation. We found that the amplitude of the early, monosynaptic component of evoked responses (the average current between 5 and 15 ms after the shock; Figure 3B) differed across experimental groups (Figure 3A; F(5,161)=3.2, p=0.009; see Methods for a description of the linear model we used). Both Sync and Async multisensory groups had larger early synaptic currents than the Control group (Tukey p=0.03 and 0.04; Cohen d=0.92 and 0.73 on log-transformed data respectively). The amplitude of late synaptic currents produced by recurrent network activation (15-145 ms after the stimulus) did not differ across groups.

**Figure 3.**
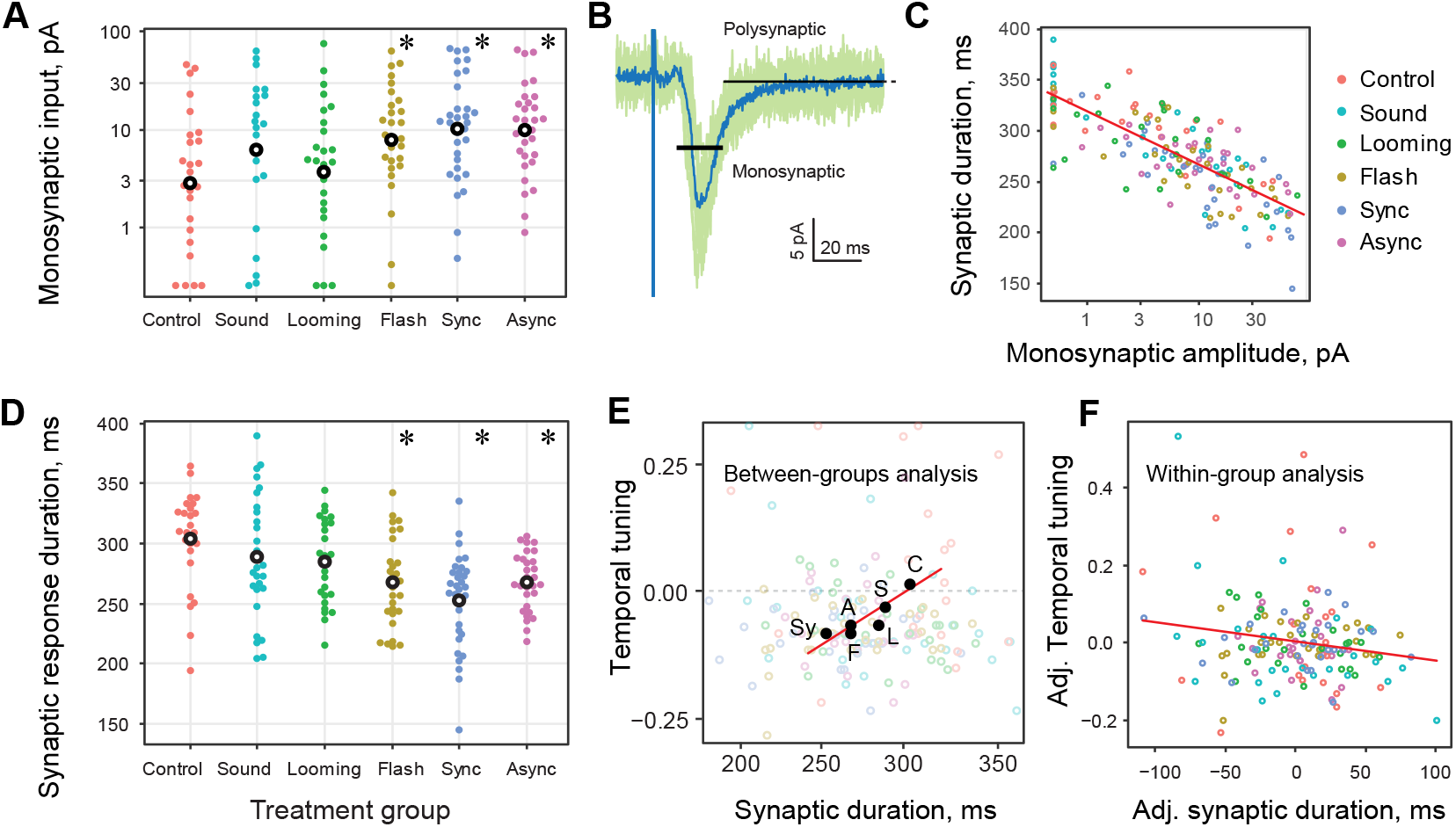
Changes in synaptic transmission, and co-tuning of synaptic and intrinsic neuronal properties. (**A**). Amplitudes of early monosynaptic inputs to tectal neurons in all experimental groups (in log scale, outliers brought within the axes limits, stars show t-test p<0.05 compared to control). (**B**). A sample synaptic recording, showing all traces for one cell (green) and an average trace (blue). The black bars show the areas at which early monosynaptic and late polysynaptic currents were measured; the vertical position of each bar represents the respective average current. The second, longer bar does not completely fit within the figure at this scale. (**C**). Synaptic current duration (vertical axis) was mostly defined by the amplitude of early monosynaptic inputs (horizontal axis). (**D**). Synaptic current durations were different between experimental groups (see text; stars show t-test p<0.05 compared to control). (**E**). Across experiments, average temporal tuning in each group positively correlated with the average durations of synaptic currents they received. (**F**). Within experimental groups, temporal tuning of individual neurons negatively correlated with the duration of synaptic currents they received. Axes show within-group deviations of temporal tuning and synaptic current duration from respective averages for each group.

To better match and compare temporal properties of synaptic inputs to those of intrinsic tuning, we calculated average “synaptic current duration” for each cell as a temporal “center of mass” of currents within the first 700 ms after optic chiasm stimulation (see Methods). Neurons with different contribution of early and late synaptic responses naturally had different synaptic current duration: cells with strong monosynaptic inputs had shorter currents, while polysynaptic activity made synaptic currents longer (Figure 3C; p=2e-16, r= −0.78, n=168). The synaptic current duration was different across treatment groups (Figure 3D; F(5,163)=6.3, p=2e-5). Cells in Flash, Sync, and Async groups all received shorter synaptic inputs than Control cells (Tukey p<0.05, mean duration of 267±36, 253±39, 268±24, and 304±40 ms, respectively). This shows that prolonged sensory activation with short, frequent stimuli reshaped synaptic transmission in the tectum, making it faster through selective potentiation of visual inputs from the eye, compared to recurrent inputs within the tectum, that remained unchanged. In this study, we did not probe changes in acoustic and mechanosensory projections from the hindbrain.

### Co-tuning of synaptic and intrinsic properties

To see whether intrinsic and synaptic temporal properties of tectal cells coordinated with each other, we compared intrinsic temporal tuning of every neuron (that is, whether it preferred longer or shorter simulated synaptic inputs in dynamic clamp experiments) to the actual duration of synaptic inputs it received, assessed by optic chiasm stimulation. We found that, *on average*, in animals exposed to stronger sensory stimuli, neurons were more responsive to shorter synaptic inputs, and also received shorter synaptic currents, leading to a positive correlation between these values across experimental groups (Figure 3E; r=0.89, p=0.02, n=5).

In contrast, *within* experimental groups, cells that preferred shorter synaptic inputs in dynamic clamp tended to receive longer actual synaptic currents, and vice versa (F(1,145)=4.9, p=0.03). The deviations of neuronal properties from their respective group averages are shown in Figure 3F. Cells that received shorter synaptic currents, compared to other cells, tended to be more responsive to longer synaptic inputs (adjusted r= −0.19, p=0.02, n=151). This suggests that during normal brain development, individual neurons tended to tune their intrinsic properties away from the typical statistics of their inputs, enhancing responses to unusual patterns of synaptic activation. This tuning to “unusual stimuli” fits into the narrative of information transfer maximization (Stemmler and Koch, 1999; Brenner et al., 2000) and network criticality (Rubinov et al., 2011), wherein every element of a network tries to maximize its influence on the overall computation. It also means that the correlation of intrinsic and synaptic properties had opposite signs between-groups (Figure 3E; positive) and within-groups (Figure 3F; negative), which is a textbook case of a so-called “Simpson’s paradox” in data analysis.

For amplitude tuning, the interaction between synaptic and intrinsic parameters of tectal cells was inconclusive. The amplitude of early synaptic responses and intrinsic amplitude tuning formally correlated on a full dataset (p=0.03, r= −0.17, n=151), but the correlation disappeared (p>0.05) when the highly non-normal amplitude data was log-transformed, or when 4 extreme values (out of 135 total) were removed. When analyzed separately, the between-groups and within-groups correlations were also insignificant.

### The mechanisms behind temporal intrinsic plasticity

Knowing that tectal neurons can tune to inputs of different temporal dynamics, we then tried to identify the cellular mechanisms underlying this tuning. For each cell, we used a sequence of voltage steps (Figure 4A) to activate Na and K conductances, and quantified ionic current amplitudes and activation potentials (Figure 4B) as it was done in earlier studies (Ciarleglio et al., 2015). Together with cell membrane resistance (Rm) and capacitance (Cm) it gave us eight intrinsic parameters for every cell: peak amplitudes for sodium current, early (transient) potassium current, and late (stable) potassium current (INa, IKt, IKs respectively), and activation potentials for these three currents (VNa, VKt, and VKs).

**Figure 4.**
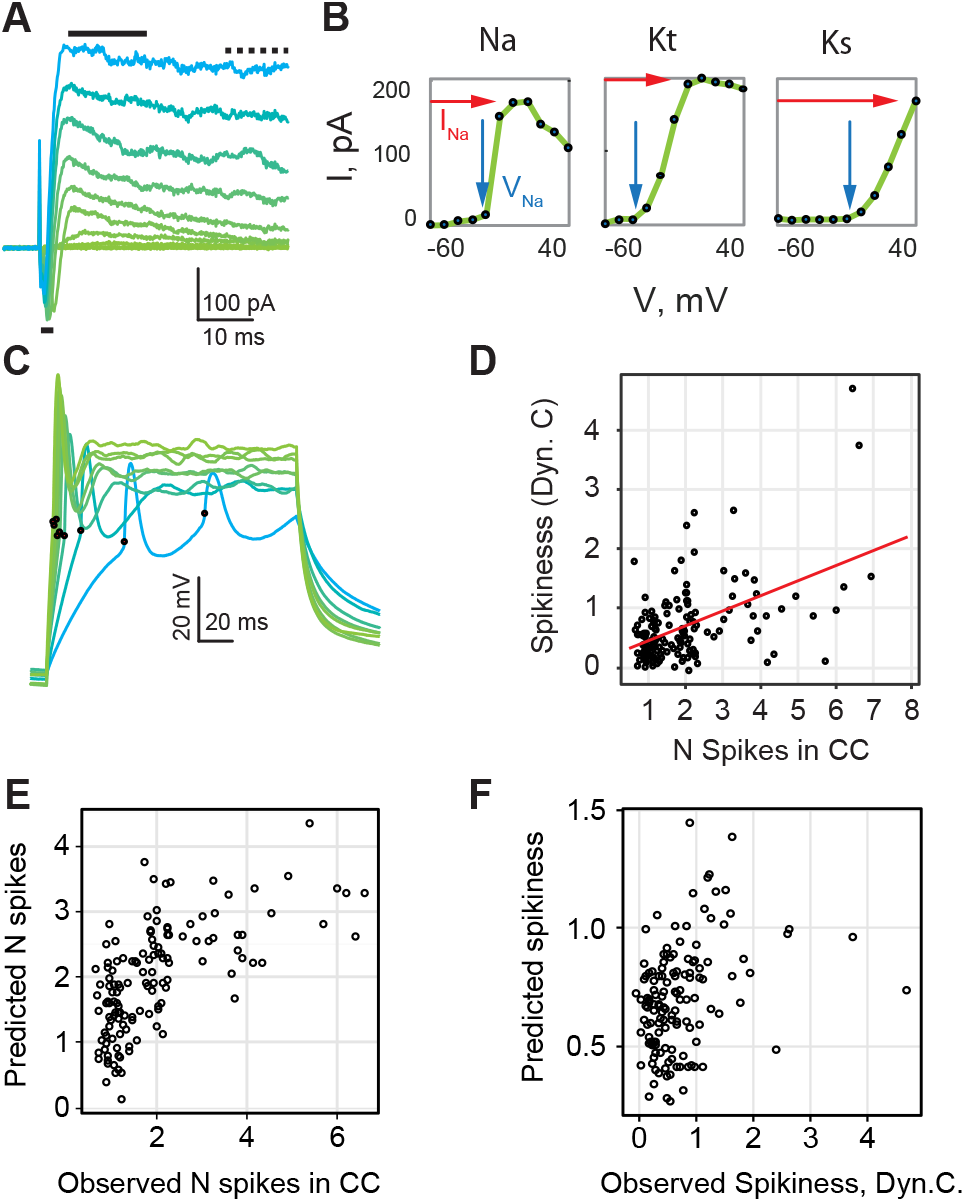
Electrophysiological properties of individual neurons compared to their spiking in current and dynamic clamp experiments. (**A**). A set of curves from one voltage step experiment; black bars show the areas used to average Na (bottom) and transient K (top) currents (see Methods). (**B**). Processing of ionic currents data with IV-curves translated into two parameters (threshold potential and peak current) for each ionic conductance. (**C**). Sample data from one current clamp experiment; spikes are marked with black dots. (**D**). Estimations of excitability from current clamp experiments (horizontal axis) and dynamic clamp experiments (vertical axis) correlate. (**E**). The number of spikes registered in current clamp mode: values predicted from a linear model plotted against observed values. The model works reasonably well (61% of variance explained). (**F**). Similar comparison for the dynamic clamp experiments: the model has very low predictive value (13% of variance explained). Here and in E, both values are adjusted for position.

We ran a stepwise generalized linear model selection analysis (R package stepAIC, Venables and Ripley 2013) to explain the intrinsic tuning of cells recorded in all experimental groups through these eight variables. We found that the average spikiness (after adjustment for cell position within the tectum; see below) was best described by a combination of sodium peak current (INa) and membrane resistance (Rm) variables, but these variables explained only 8% and 2% of cell-to-cell variance respectively (F(1,130)=11.3, and F(1,130)=4.5; Figure 3E). Together, all eight cellular parameters described only 13% of variance in average spikiness. The temporal tuning value was best explained by sodium current activation potential and membrane resistance (VNa: 7%, F(1,147)=10.7; Rm: 2%, F(1,147)=3.1), with all eight variables explaining only 11% of total variance. For amplitude tuning, the proposed best model included peak sodium current (INa: 6%, F(1,163)=10.0) and sodium activation potential (VNa: 2%, F(1,1,163)=3.5), with all eight variables accounting for 10% of variance. This very low total explained variance indicates that while ionic currents and their activation potentials clearly affected intrinsic tuning of tectal cells, most cell-to-cell variability in intrinsic phenotypes stemmed from some other properties that we did not measure in voltage-clamp experiments. In agreement with this assessment, the effect of experimental group on either mean spiking, temporal, or amplitude tuning curves remained significant even after compensating for all 8 intrinsic properties (sequential sum of squares analysis of variance p=0.03, 0.001, and 0.003 respectively).

### A comparison between dynamic clamp and current clamp experiments

The inability to predict spiking of tectal neurons from their isolated electrophysiological properties was unexpected, and stood in a seeming contradiction with our previous study (Ciarleglio et al., 2015). Fortunately, in the current study, we recorded spiking traces in response to “classic” current steps (Figure 4C), which allowed a direct comparison between the results of current clamp and dynamic clamp protocols. Across all cells, the maximal number of spikes observed during current step injections correlated with the average number of spikes in dynamic clamp experiments (Figure 4D; r=0.46, p=2e-9, n=152). In agreement with (Ciarleglio et al., 2015), spiking in current clamp experiments correlated with peak sodium (INa: r=0.42, p=2e-8, n=152) and stable potassium currents (IKs: r=0.39, p=2e-7), as well as activation potential for sodium current (VNa: r=0.24, p=0.02). Together, the 8 intrinsic variables described above (Rm, Cm, three peak currents, and three activation potentials) explained 61% of cell-to-cell variability in the maximal number of spikes from current clamp experiments (Figure 4E), which is comparable to 49% reported in (Ciarleglio et al., 2015), and noticeably higher than 13% for dynamic clamp experiments (Figure 4F).

We can therefore conclude that our set of eight cellular parameters can much better predict spiking during current step injections (61% of variance), compared to more physiologically realistic dynamic clamp experiments (13% of variance). This suggests the existence of intrinsic properties that strongly affect spiking in dynamic clamp experiments, but are inaccessible through simple voltage step protocols, such as the amount of voltage-gated channels inactivation (Zbili et al. 2019; see Discussion). Our hypothesis is also indirectly supported by the observation that only one out of eight cellular parameters was significantly (p<0.05) different across treatment groups (NaV: F(5,175)=3.7, p=0.003), and that the number of spikes detected in current clamp experiments did not differ across experimental groups (F(5,165)=0.8, p=0.6).

### Effects of position within the tectum

In all analyses presented above, we adjusted cell properties for rostro-caudal and medio-lateral position of each cell within the tectum, as in tadpoles both intrinsic (Hamodi and Pratt, 2014) and synaptic properties (Wu et al., 1996; Khakhalin and Aizenman, 2012) differ between older (rostro-medial) and younger (caudal, lateral) parts of the developing tectum. Most cell properties we measured in this study correlated (p<0.05 after adjustment for treatment group differences) with either medial or rostral position within the tectum (medial: membrane capacitance r= −0.17, membrane resistance r=0.25, sodium current activation potential r= −0.30, stable potassium current r= −0.16, early synaptic amplitude r= −0.22, synaptic current duration r=0.29; rostral: peak sodium current r=0.18, late synaptic amplitude r=0.11; n between 168 and 183). Curiously, neither of the three measures of intrinsic tuning (average spikiness, temporal tuning, and amplitude tuning) correlated with position (p>0.1, n=168). This seems to suggest that while low-level properties of tectal cells depended on their developmental age, their spiking phenotypes were age-independent. This means that different cells achieved similar spiking behaviors through different combinations of underlying parameters, relying on the principle of “parameter degeneracy” (Prinz et al., 2004b; Drion et al., 2015).

## Discussion

In this study, we show that stimuli of different temporal dynamics differentially retune *Xenopus* tectal neurons, changing both their temporal and amplitude transfer functions (Figure 2). This addresses the first question of this study, about the functional scope of intrinsic plasticity in the tectum, and shows that it goes well beyond simple adjustments of neuronal excitability.

In answer to our second question about whether intrinsic and synaptic temporal properties of tectal cells are in any way coordinated, we show that they are weakly co-tuned, and moreover, that this co-tuning is modified by sensory experience (Figure 3). In contrast with earlier studies that reported increased excitability after sensory stimulation (Aizenman et al., 2003; Dong et al., 2009; Ciarleglio et al., 2015), we found that stimulation led to a suppression of spiking, and potentiation of fast monosynaptic inputs from the eye. We believe that the reason for this difference is that in earlier studies, the visual stimulation was provided with very bright, high-contrasted LEDs, which caused a suppression of synaptic inputs via a polyamine block of AMPA receptors (Aizenman et al., 2003). This suppression then triggered a “second-order” homeostatic compensation (Turrigiano, 2011; Tien and Kerschensteiner, 2018), making neurons spikier. In our current experiments, synaptic inputs were not suppressed, and so synaptic drive during sensory conditioning had to be stronger than in control (Khakhalin et al., 2014), causing a decrease in intrinsic excitability.

Trying to answer the third question, about the potential mechanism behind the intrinsic temporal tuning variability, we found that tuning of individual tectal cells could not be reliably predicted from their lower-level electrophysiological properties (Figure 4). This made our dynamic clamp results (Figure 1) very different from the results of a “classic” current clamp protocol (Figure 4C). While in dynamic clamp spiking responses were temporally tuned, affected by stimulation, and did not coordinate with low-level intrinsic parameters, spiking recorded in a standard current clamp mode did not show temporal tuning (Ciarleglio et al., 2015), was not shaped by stimulation (in this study), yet was well coordinated with intrinsic parameters.

This difference between neuronal responses in the dynamic clamp and fixed current clamp experiments may be interpreted in two ways. A pessimistic take would be to argue that the dynamic clamp in the soma offered a bad approximation of peripheral synaptic inputs, as space clamp errors are known to be more pronounced for fast voltage waveforms than for constant current injections (Spruston et al., 1993; Prinz et al., 2004a). We however find this hypothesis unlikely, as our dynamic clamp responses were consistently different in animals with different sensory history, suggesting that we have captured some important aspects of intrinsic tuning diversity, even if our estimations were biased.

Another, and in our opinion more likely explanation, is that intrinsic excitability of tectal cells is shaped by mechanisms that are not easily accessible by slow voltage- and current-clamp protocols, such as changes in voltage-gated channel inactivation (Azouz and Gray, 2000), and axon initial segment decoupling (Grubb and Burrone, 2010; Kuba et al., 2010; Kole and Stuart, 2012). Let us address these two potential mechanisms separately.

First, while tectal neurons lack “true” resonant currents, such as Ih currents (Ciarleglio et al., 2015), much of temporal tuning can be achieved through the adjustment of sodium and potassium current inactivation (Azouz and Gray, 2000; Fontaine et al., 2014), either via channel phosphorylation, or through the expression of channels with different inactivation dynamics (Frank and Catterall, 2003; Goldwyn et al., 2018). Crucially however, if these channels are located far from the soma, a prolonged current injection would quickly inactivate them, obscuring any possible interplay between action potential shape and sodium channel recovery that are critical for burst firing (Popovic et al., 2011; Kole and Stuart, 2012; Fontaine et al., 2014). These effects however would be fully at play during responses to faster, and more biophysically realistic dynamic clamp experiments (Clay et al., 2012; Zbili et al., 2019).

Second, we know that some subcellular compartments, such as the axon initial segment or excitable dendrites, have a disproportionally strong influence on neuronal tuning (Jarvis et al., 2018; Moldwin and Segev, 2019). Even a minor change in electrical coupling between these compartments, either because of structural changes in cell morphology (Grubb and Burrone, 2010; Kuba et al., 2010; Leterrier, 2018), or due to modulation of channels at key integrative positions within the dendritic tree (Murakoshi et al., 1997), may lead to drastic changes in neuronal responses. And yet, the channels at these key positions may constitute only a small fraction of all voltage-gated channels in the cell, making them practically “invisible” in classic current clamp experiments (Kole et al., 2007; Hamada et al., 2016).

The hypothesis that tuning of tectal neurons may rely on subtle reorganization of their dendritic trees, or axons branching from these trees (Lazar, 1973), is also particularly exciting in view of the unexplained finding from our earlier study, in which cell membrane capacitance (Cm) was found to decrease after strong sensory stimulation (Ciarleglio et al. 2015, Figure 7D). Cell membrane capacitance is often thought to be relatively immutable, and is even used to estimate cell size. For this reason, the change in electrical coupling between different compartments of a cell may be one the few plausible explanations for the rapid modulation of this parameter in response to sensory stimulation.

The best way to differentiate between these two hypotheses would be to explicitly measure sodium and potassium current inactivation (Zbili et al., 2019) in tadpoles exposed to different types of sensory stimulation. One could also use pharmacological interventions to provide negative and positive control groups for this study. It would be even better to follow it up with immunostaining of cleared tectal preparations, and look whether the distribution of sodium and potassium channels is changed after sensory stimulation. Finally, all these observations can be validated in a biophysical computational model of a tectal neuron.

Our findings also lead to several verifiable predictions. As rapid inactivation of spiking in tectal neurons plays a role in collision detection (Khakhalin et al., 2014; Jang et al., 2016), a change in temporal tuning should affect loom detection, which can be verified experimentally. We predict that instantaneous flashes that suppress responses to slow inputs would selectively disrupt slow collision avoidance (Khakhalin et al., 2014). The changes in intrinsic temporal tuning would also reshape tectal connectivity, as fast-inactivating cells would not support short recurrent loops within the network, thus indirectly promoting long-ranged polysynaptic connectivity (Fiete et al., 2010; Clopath et al., 2010; Khakhalin, 2019). Finally, based on the multisensory phenomena reported in this paper, we predict that even though multisensory stimulation tends to increase tectal responses in vitro (Felch et al., 2016; Truszkowski et al., 2017), it may actually reduce peak activation in vivo.

To sum up, we present a novel case of temporal selectivity in non-oscillatory neurons in a sensory network, and demonstrate that intrinsic temporal tuning of neural cells correlates with their synaptic properties, and is modified by sensory experiences. We also argue that any cell with a sufficiently complex morphology could be able to tune its intrinsic temporal selectivity in ways we describe in this paper. Crucially, we expect that in many cases this tuning would not be noticeable in experiments that use standard voltage- and current-clamp protocols, yet can be easily uncovered with a dynamic clamp technique. It would therefore be interesting to see whether our results will replicate in other sensory systems.

## Acknowledgements

We would like to thank Carlos Aizenman (Brown University), Justin Hulbert (Bard College), and Kara Pratt (University of Wyoming) for their feedback on drafts of this paper. Part of this work was supported by the Bard Summer Research Institute (BSRI) program.

The authors have no conflicts of interest to disclose.

## Author Contributions

S.E.B.: Conception and design, acquisition of data, analysis and interpretation of data, drafting and revising the article.

A.S.Kh.: Conception and design, analysis and interpretation of data, figure preparation, drafting and revising the article.

## Materials and Methods

### Housing and sensory conditioning

All experimental protocols were in accordance with Bard College Institutional Animal Care and Use Committee (IACUC), and National Institutes of Health (NIH) guidelines. Animals were purchased from Nasco (Fort Atkinson, WI, USA) at developmental stages 44-47, and raised to stages 48-49 on a 12/12 h light/dark cycle at 18 °C.

In the beginning of each experiment, a tadpole was put in a Petri dish (diameter of 10 cm) filled with 1-1.2 cm of tadpole rearing medium, placed on top of a CRT monitor with two speakers connected to the Petri dish with short wooden struts (James et al., 2015; Truszkowski et al., 2017), and kept there for 4 hours. The tadpole was visually isolated from the rest of the room with a cardboard box surrounding the apparatus. For Control and Sound groups, the monitor was on, but showed a uniform 50% gray background. For Flash, Sync, and Async groups the screen showed a black-and-white checkerboard pattern, with each square in the pattern being 14 mm wide; this pattern flipped (inverted) every 1 second. For the Looming group, the inversion of the pattern was not instantaneous, but lasted for one second, with old black squares linearly shrinking into white background, and new black squares appearing and linearly expanding in the middle of each white square (Figure 1C). The stimulation program was written in JavaScript, using the p5.js library (McCarthy et al., 2015), and is available at http://faculty.bard.edu/~akhakhal/checker_flash_ding.html. For Sound, Sync, and Async groups a broad-spectrum sound click was delivered through the speakers every 1 second, with left and right speakers playing the same waveform, but inverted. Formally the click was generated as a 5 ms pulse of 100 Hz sine wave, but it was also distorted by the non-linearities in the system. The sound volume was calibrated to be about 2 times higher than the threshold volume, which means that it reliably evoked startle responses with about 80% success ratio, at least at the beginning for the conditioning protocol. For the Sync group, the sound clicks and the checkerboard inversions were synchronized, while for the Async group they were offset by 500 ms (half a period).

### Electrophysiology

Immediately after sensory conditioning, tadpoles were anesthetized in 0.02% tricaine methanesulfanate (MS-222). Dorsal commissures were cut, the brain was dissected out (Aizenman et al., 2003; Ciarleglio et al., 2015), and placed in the recording chamber filled with artificial cerebrospinal fluid (in mM: 115 NaCl, 4 KCl, 3 CaCl2, 3 MgCl2, 5 HEPES, 10 glucose, 10 *µ*M glycine; pH 7.2, osmolarity 255 mOsm). All chemicals were obtained from Sigma (Sigma-Aldrich, St. Louis, MO). The ventricular membrane was removed (suctioned) using a broken glass electrode. Cells were visualized with a Nikon (Tokyo, Japan) Eclipse FN1 light microscope with a 40x water immersion objective. Recordings were restricted to the middle of the tectum, as in earlier studies (Ciarleglio et al., 2015), from 25% to 53% of brain half-width medially from the lateral edge, and from 36% to 69% of tectum length rostrally from the caudal edge of the tectum (Figure 1A). Care was taken to record only from “deep” primary tectal cells (that are located superficially in our preparation), and not from MV cells (Pratt and Aizenman, 2009) or superficial layer cells (that are located deep in the tectum in our preparation) (Liu et al., 2016). Glass electrodes (1.5×0.86 mm borosilicate glass; Sutter instruments, Novato, CA) were pulled on a Sutter P-1000 puller (Sutter instruments), to a tip resistance of 8-12 MOhm. The elecrodes were filled with intracellular saline (in mM: 100 K-gluconate, 5 NaCl, 8 KCl, 1.5 MgCl2, 20 HEPES, 10 EGTA, 2 ATP, 0.3 GTP; pH 7.2, osmolarity 255 mOsm). Electrodes were placed in an Axon headstage (Molecular Devices, Sunnyvale, CA), controlled by a motorized micromanipulator (MX7600, Siskiyou, Grants Pass, OR). Whole cell patch clamp was established as usual (Ciarleglio et al., 2015), with typical final access resistance of 30 MOhm, and membrane resistance Rm of 3.2 GOhm. Signals were measured with an Axon Instruments MultiClamp 700B amplifier (Axon Instruments, Foster City, CA), filtered with a 5 kHz band-pass filter, and digitized at 10 kHz a CED Power1401-3 Digitizer (Cambridge Electronic Design; Cambridge, England). For synaptic stimulation, a bipolar stimulating electrode (Warner Instruments, Hamden, CT) was placed on the optic chiasm (Wu et al., 1996); stimuli were controlled by a CED digitizer, and were delivered by A.M.P.I. stimulus isolator (AMPI, Jerusalem, Israel).

Each neuron was subjected to a series of electrophysiological measurement protocols (see below for details), closely matching experimental protocols from (Ciarleglio et al., 2015). For each cell, we measured membrane resistance Rm and capacitance Cm in voltage clamp mode, and then (1) ran a series of voltage steps to measure ionic currents; (2) in current clamp mode, ran a series of current steps to assess cell spiking; (3) in dynamic clamp mode, subjected the cell to different conductance injections; (4) finally, if the cell was still in good health, we ran a synaptic protocol with optic chiasm stimulation. All data was processed offline using custom Matlab scripts (Mathworks, Natick, MA), and analyzed in R. In total, we recorded from 188 neurons in 35 tadpoles; of these, 159 cells had readings from all 4 protocols, while 12 lacked synaptic recordings; these 12 cells were scattered across all 6 experimental groups. After the recording was over, the position of each recorded cell was visualized with a 10x microscope, marked on a screen, measured in medial and rostral directions relative to the most latero-caudal point of the tectum, and converted into a percentage (Hamodi and Pratt, 2014).

### Voltage steps protocol

The baseline membrane potential was set at –60 mV (in this manuscript, the voltages are not adjusted for junction potential, which is expected to be equal to –12 mV for this combination of external and internal solutions). After Cm and Rm were measured with a standard seal test, cells were subjected to 11 voltage steps (square pulses), each 500 ms long, and 10 mV higher than the previous one, with 500 ms of baseline voltage between the steps. Each trial also contained a 50 ms long test pre-step of –10 mV relative to the baseline. During analysis, we averaged transition currents evoked by the leading and trailing edges of the pre-step, then scaled and subtracted them from the current responses to the main step. For remaining active currents, we measured average currents during a 0.4-2.7 ms range after the step (Na current), 5.7-19.7 ms after the step (Kt, or transient potassium current), and 430-490 ms (Ks, or stable potassium current). This approach is standard for recordings from the *Xenopus* tectum, as ionic currents are slow enough to be separated temporally (Aizenman et al., 2003). The ionic conductances were quantified as is (Ciarleglio et al., 2015). For each cell, the values of current as a function of voltage were fit with an empirical parametric equation:

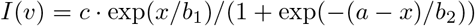

for Na and Kt currents (sigmoid, followed by exponential decay, inactivating), and a different equation:

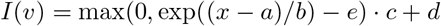

for Ks current (a shifted piece of exponentially increasing curve with its lower part cut off; not inactivating). For equations with inactivation, we used its *I*_*max*_ value as a measure of amplitude, and *v*_*a*_ on the rising front such that *I*(*v*_*a*_) = *I*_*max*_/2 calculated the threshold potential. For curves without inactivation (Ks) we used *I*_*max*_ and the first non-zero point (*v*_*a*_ = *a* + log(*e*) *⋅ b*) for the same purpose.

As a preliminary verification of our results, we compared the overall structure of our new dataset with the dataset from the 2015 study (Ciarleglio et al., 2015). The eight cellular parameters described above showed a similar pattern of coordination in both datasets: 23 pairwise correlations out of 35 total were significant (p<0.05) in this dataset, compared to 21 out of 35 in the 2015 study. The average absolute value of correlation coefficient was r=0.38 in this study, compared to r=0.32 in the 2015 study. This suggests that the datasets are similar and representative of true internal variability in the tectum.

### Current steps protocol

For the current steps protocol, we switched each cell to current clamp mode, and adjusted the stable holding current to bring the resting membrane potential to about –60 mV. We then subjected the cell to 10 current pulses, each 150 ms in duration, delivered every 1 s, such that currents ranged from 0 to 180 pA in 20 pA increments. Cells that did not produce at least one spike in this experiment were considered not-excitable, and were not included in the dataset. The largest number of spikes produced in response to a single current injection was estimated offline, manually, using a custom Matlab data browser that blinded the researcher to the identity of the cell. As a control, spikes were also detected automatically, using the filtering and thresholding approach that was used in (Ciarleglio et al., 2015); in 78% of cells both manual and automated estimations matched, in remaining 22% of cells the mismatch was either due to artifacts on the rising front being auto-detected as spikes, or due to spike broadening that fell below the threshold for the adaptive filter. The number of cells in which manual spike detection disagreed with automated detection did not differ across groups (6.1*±*1.7; p=0.5, exact Fisher test).

### Dynamic clamp protocol

For dynamic clamp experiments, each cell was held at −50 mV baseline potential, and was stimulated with 5 repetitions of 12 different “conductance injections”. Conductance curves were generated with a formula 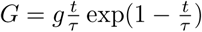, known as “alpha synapse”, where *g* and *τ* are conductance and decay parameters respectively (Destexhe et al., 1994). We used four different values of *τ*, to represent four typical patterns of synaptic activation: 20 ms, corresponding to a total curve length (decay to 10% of the peak value) of about 100 ms, to approximate short, monosynaptic inputs (Ciarleglio et al., 2015); 40 ms, corresponding to a total curve length of about 200 ms, as for a typical in-vitro stimulus with polysynaptic activation (Xu et al., 2011); 100 ms, to mimic in-vivo inputs to the tectum in response to abrupt disappearance of light (“dark-flash”) (Khakhalin et al., 2014); and 200 ms, to mimic retinal inputs in response to a 1 second-long linear looming stimulus (Khakhalin et al., 2014). Actual decay times to 10% of peak amplitude were 98, 196, 489, and 978 ms respectively. The value of *g* was adjusted so that conductance curves peaked at 3 target conductances of 0.2, 0.5, and 1 nS. With the cell clamped at –50 mV, these conductances would have induced currents that peaked at 10, 25, and 50 pA respectively, matching the range of peak synaptic currents observed in (Xu et al., 2011; Khakhalin et al., 2014; Ciarleglio et al., 2015). Conductance curves were always presented in the order from the shortest to the longest, and this sequence was not randomized.

For each cell, for each of 60 trials (5 repetitions of 12 conductances), spikes were counted manually, blindly, and independently by both authors, using a custom Matlab data browser script. There was a 98.5% agreement between spike number estimations on a trace-by-trace basis. All cases of disagreement (usually ±1 spike) were due to later action potentials becoming broader and smaller in amplitude, which made them ambiguous. We ran sensitivity analyses of main effects reported in this paper separately on both estimations, and got qualitatively identical results. Numerically, we went with consensus numbers that in each case followed the higher estimation for the number of spikes.

To quantify the “shape” of spiking responses to conductances of different duration (temporal tuning), we encoded curve duration as an ordinal value (from 1 to 4) for every cell, fit the spike data as a function of response duration with a quadratic formula (*y* = *ax*^2^ + *bx* + *c*), and used the quadratic coefficient *a* as the measure of response non-linearity. While the numerical values of this dimensionless coefficient are not easily interpretable, it captures the shape of the response curve well, and allows for easy comparisons between cells (Figure 2A). The case of *a* = 0 corresponds to spiking output linearly increasing with duration increase; *a* > 0 means supralinear preference for long conductances (curving up); about –0.25 < *a* < 0 corresponds to a plateau-shaped curves, while *a* < –0.25 would mean heavy spike inactivation for longer conductance injections.

### Synaptic recordings

For synaptic recordings, we switched cells back to voltage clamp mode, and held the membrane potential at –45 mV to isolate excitatory synaptic currents. Optic chiasm shocks were delivered 10 times, every 20 s, with a stimulation strength between 0.05 and 0.4 mA, and with a pulse length of 0.2 us. In each experiment, we would first find stimulation strength that evoked consistent synaptic responses in the first cell we patched, then increased it by 20% and kept it constant for all cells recorded from that brain. Recordings were processed offline; for each trial we used the average current between 5 and 15 ms as a measure of monosynaptic response amplitude, and current between 15 and 145 ms as a measure of polysynaptic response amplitude (Ciarleglio et al., 2015). The weighted duration of synaptic responses was calculated as the “center of mass” under the first 700 ms of the curve:

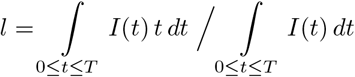

### Statistics and reporting

To analyze the numbers of spikes observed in dynamic clamp experiments (Figure 1), we first averaged the number of generated spikes for each combination of conductance curve duration and amplitude, 12 values for each of 5 protocol repetitions for every cell. Then we used sequential sum of squares analysis of variance with repeated measures. Both different conductance curve amplitudes and durations were represented as ordinal values (1 to 3 for amplitude, 1 to 4 for duration). Differences between experimental groups were assessed as interactions between these ordinal values and the factor variable encoding the experimental group, as we were interested in response shapes (reflected by interactions) rather than average values of spikiness (reflected by independent terms). Cell ids were included in the analysis as a fixed factor for repeated measures analysis of variance (also equivalent to analysis of variance with blocking). To verify the validity of this approach, we also ran a maximal likelihood mixed-effects model with type III interaction terms, as implemented in R package “lmer”, with “lmerTest” extension to get access to Satterthwaite degrees of freedom and p-value estimations (Kuznetsova et al., 2017). The results of both methods were numerically similar, and supported same conclusions.

For the comparison of summative descriptions of tuning, and other electrophysiological cell parameters between experimental groups, we report p-values of fixed effect sequential sum of squares linear model (ancova), in which rostral and medial coordinates of each cell within the tectum are included as covariates, and experimental group is used as the main factor. All comparisons and correlations between cell parameters are performed on values adjusted for cell position within the tectum. Position adjustment was based on a two-way linear regression model without interaction. For five variables that were distributed extremely non-normally, this adjustment for position was performed on transformed values (original values were transformed to normally distributed proxy values, linearly adjusted, and then transformed back): for early and late mean synaptic amplitudes we used a transformation *a*′ = *log*(1 – *a*); for the variability of synaptic amplitudes *s*′ = *log*(1 + *s*); and for temporal tuning 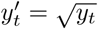. Where appropriate, we performed the analysis with and without extreme outliers, and reported the difference. All analyses presented in the paper were also verified in mixed model analyses, with animal id included as a random factor; the results of these mixed model analyses were similar to that of a fixed model, and are not reported.

## References

Aizenman, C. D., Akerman, C. J., Jensen, K. R., and Cline, H. T. (2003). Visually driven regulation of intrinsic neuronal excitability improves stimulus detection in vivo. Neuron, 39(5):831–842.

Aizenman, C. D. and Cline, H. T. (2007). Enhanced visual activity in vivo forms nascent synapses in the developing retinotectal projection. Journal of neurophysiology, 97(4):2949–2957.

Azouz, R. and Gray, C. M. (2000). Dynamic spike threshold reveals a mechanism for synaptic coincidence detection in cortical neurons in vivo. Proceedings of the National Academy of Sciences, 97(14):8110–8115.

Bassett, D. S. and Sporns, O. (2017). Network neuroscience. Nature neuroscience, 20(3):353.

Beatty, J. A., Song, S. C., and Wilson, C. J. (2014). Cell-type-specific resonances shape the responses of striatal neurons to synaptic input. Journal of neurophysiology, 113(3):688–700.

Bianchi, D., Marasco, A., Limongiello, A., Marchetti, C., Marie, H., Tirozzi, B., and Migliore, M. (2012). On the mechanisms underlying the depolarization block in the spiking dynamics of ca1 pyramidal neurons. Journal of computational neuroscience, 33(2):207–225.

Branco, T., Clark, B. A., and Häusser, M. (2010). Dendritic discrimination of temporal input sequences in cortical neurons. Science, 329(5999):1671–1675.

Brenner, N., Bialek, W., and Van Steveninck, R. d. R. (2000). Adaptive rescaling maximizes information transmission. Neuron, 26(3):695–702.

Ciarleglio, C. M., Khakhalin, A. S., Wang, A. F., Constantino, A. C., Yip, S. P., and Aizenman, C. D. (2015). Multivariate analysis of electrophysiological diversity of xenopus visual neurons during development and plasticity. Elife, 4.

Clay, J. R., Forger, D. B., and Paydarfar, D. (2012). Ionic mechanism underlying optimal stimuli for neuronal excitation: role of na+ channel inactivation. PloS one, 7(9):e45983.

Clopath, C., Büsing, L., Vasilaki, E., and Gerstner, W. (2010). Connectivity reflects coding: a model of voltage-based stdp with homeostasis. Nature neuroscience, 13(3):344.

Deeg, K. E. and Aizenman, C. D. (2011). Sensory modality–specific homeostatic plasticity in the developing optic tectum. Nature neuroscience, 14(5):548.

Deeg, K. E., Sears, I. B., and Aizenman, C. D. (2009). Development of multisensory convergence in the xenopus optic tectum. Journal of Neurophysiology, 102(6):3392–3404.

Destexhe, A., Mainen, Z. F., and Sejnowski, T. J. (1994). Synthesis of models for excitable membranes, synaptic transmission and neuromodulation using a common kinetic formalism. Journal of computational neuroscience, 1(3):195–230.

Dong, W., Lee, R. H., Xu, H., Yang, S., Pratt, K. G., Cao, V., Song, Y.-K., Nurmikko, A., and Aizenman, C. D. (2009). Visual avoidance in xenopus tadpoles is correlated with the maturation of visual responses in the optic tectum. Journal of neurophysiology, 101(2):803–815.

Drion, G., O’Leary, T., and Marder, E. (2015). Ion channel degeneracy enables robust and tunable neuronal firing rates. Proceedings of the National Academy of Sciences, 112(38):E5361–E5370.

Evans, M. D., Dumitrescu, A. S., Kruijssen, D. L., Taylor, S. E., and Grubb, M. S. (2015). Rapid modulation of axon initial segment length influences repetitive spike firing. Cell reports, 13(6):1233–1245.

Felch, D. L., Khakhalin, A. S., and Aizenman, C. D. (2016). Multisensory integration in the developing tectum is constrained by the balance of excitation and inhibition. Elife, 5.

Fiete, I. R., Senn, W., Wang, C. Z., and Hahnloser, R. H. (2010). Spike-time-dependent plasticity and heterosynaptic competition organize networks to produce long scale-free sequences of neural activity. Neuron, 65(4):563–576.

Fontaine, B., Peña, J. L., and Brette, R. (2014). Spike-threshold adaptation predicted by membrane potential dynamics in vivo. PLoS computational biology, 10(4):e1003560.

Frank, H. Y. and Catterall, W. A. (2003). Overview of the voltage-gated sodium channel family. Genome biology, 4(3):207.

Goldwyn, J. H., Slabe, B. R., Travers, J. B., and Terman, D. (2018). Gain control with a-type potassium current: Ia as a switch between divisive and subtractive inhibition. arXiv preprint arXiv:1802.04794.

Grubb, M. S. and Burrone, J. (2010). Activity-dependent relocation of the axon initial segment fine-tunes neuronal excitability. Nature, 465(7301):1070.

Hamada, M. S., Goethals, S., de Vries, S. I., Brette, R., and Kole, M. H. (2016). Covariation of axon initial segment location and dendritic tree normalizes the somatic action potential. Proceedings of the National Academy of Sciences, 113(51):14841–14846.

Hamodi, A. S., Liu, Z., and Pratt, K. G. (2016). An nmda receptor-dependent mechanism for subcellular segregation of sensory inputs in the tadpole optic tectum. eLife, 5.

Hamodi, A. S. and Pratt, K. G. (2014). Region-specific regulation of voltage-gated intrinsic currents in the developing optic tectum of the xenopus tadpole. Journal of neurophysiology, 112(7):1644–1655.

Hildebrand, D. G. C., Cicconet, M., Torres, R. M., Choi, W., Quan, T. M., Moon, J., Wetzel, A. W., Champion, A. S., Graham, B. J., Randlett, O., et al. (2017). Whole-brain serial-section electron microscopy in larval zebrafish. Nature, 545(7654):345.

Hiramoto, M. and Cline, H. T. (2009). Convergence of multisensory inputs in xenopus tadpole tectum. Developmental neurobiology, 69(14):959–971.

James, E. J., Gu, J., Ramirez-Vizcarrondo, C. M., Hasan, M., Truszkowski, T. L., Tan, Y., Oupravanh, P. M., Khakhalin, A. S., and Aizenman, C. D. (2015). Valproate-induced neurodevelopmental deficits in xenopus laevis tadpoles. Journal of Neuroscience, 35(7):3218–3229.

Jang, E. V., Ramirez-Vizcarrondo, C., Aizenman, C. D., and Khakhalin, A. S. (2016). Emergence of selectivity to looming stimuli in a spiking network model of the optic tectum. Frontiers in neural circuits, 10:95.

Jarvis, S., Nikolic, K., and Schultz, S. R. (2018). Neuronal gain modulability is determined by dendritic morphology: A computational optogenetic study. PLoS computational biology, 14(3):e1006027.

Khakhalin, A. (2019). Graph analysis of looming-selective networks in the tectum, and its replication in a simple computational model. BioRxiv, page 589887.

Khakhalin, A. S. and Aizenman, C. D. (2012). Gabaergic transmission and chloride equilibrium potential are not modulated by pyruvate in the developing optic tectum of xenopus laevis tadpoles. PLoS One, 7(4):e34446.

Khakhalin, A. S., Koren, D., Gu, J., Xu, H., and Aizenman, C. D. (2014). Excitation and inhibition in recurrent networks mediate collision avoidance in xenopus tadpoles. European Journal of Neuroscience, 40(6):2948–2962.

Kole, M. H., Letzkus, J. J., and Stuart, G. J. (2007). Axon initial segment kv1 channels control axonal action potential waveform and synaptic efficacy. Neuron, 55(4):633–647.

Kole, M. H. and Stuart, G. J. (2012). Signal processing in the axon initial segment. Neuron, 73(2):235–247.

Kuba, H., Oichi, Y., and Ohmori, H. (2010). Presynaptic activity regulates na+ channel distribution at the axon initial segment. Nature, 465(7301):1075.

Kuznetsova, A., Brockhoff, P. B., and Christensen, R. H. (2017). lmertest package: Tests in linear mixed effects models. Journal of Statistical Software, 82(13):1–26.

Lazar, G. (1973). The development of the optic tectum in xenopus laevis: a golgi study. Journal of Anatomy, 116(Pt 3):347.

Leterrier, C. (2018). The axon initial segment: an updated viewpoint. Journal of Neuroscience, 38(9):2135–2145.

Liu, Z., Ciarleglio, C. M., Hamodi, A. S., Aizenman, C. D., and Pratt, K. G. (2016). A population of gap junction-coupled neurons drives recurrent network activity in a developing visual circuit. Journal of neurophysiology, 115(3):1477–1486.

Lukoševičius, M. and Jaeger, H. (2009). Reservoir computing approaches to recurrent neural network training. Computer Science Review, 3(3):127–149.

Ma, M. and Koester, J. (1996). The role of k+ currents in frequency-dependent spike broadening in aplysia r20 neurons: a dynamic-clamp analysis. Journal of Neuroscience, 16(13):4089–4101.

Marcelin, B., Chauvière, L., Becker, A., Migliore, M., Esclapez, M., and Bernard, C. (2009). H channel-dependent deficit of theta oscillation resonance and phase shift in temporal lobe epilepsy. Neurobiology of disease, 33(3):436–447.

Marder, E. and Taylor, A. L. (2011). Multiple models to capture the variability in biological neurons and networks. Nature neuroscience, 14(2):133.

McCarthy, L., Reas, C., and Fry, B. (2015). Getting Started with P5. js: Making Interactive Graphics in JavaScript and Processing. Maker Media, Inc.

Moldwin, T. and Segev, I. (2019). Perceptron learning and classification in a modeled cortical pyramidal cell. bioRxiv.

Murakoshi, H., Shi, G., Scannevin, R. H., and Trimmer, J. S. (1997). Phosphorylation of the kv2. 1 k+ channel alters voltage-dependent activation. Molecular pharmacology, 52(5):821–828.

Narayanan, R. and Johnston, D. (2008). The h channel mediates location dependence and plasticity of intrinsic phase response in rat hippocampal neurons. Journal of Neuroscience, 28(22):5846–5860.

Ohtsuki, G. and Hansel, C. (2018). Synaptic potential and plasticity of an sk2 channel gate regulate spike burst activity in cerebellar purkinje cells. iScience, 1:49–54.

O’Leary, T., Williams, A. H., Caplan, J. S., and Marder, E. (2013). Correlations in ion channel expression emerge from homeostatic tuning rules. Proceedings of the National Academy of Sciences, 110(28):E2645–E2654.

Picton, L. D., Sillar, K. T., and Zhang, H.-Y. (2018). Control of xenopus tadpole locomotion via selective expression of ih in excitatory interneurons. Current Biology.

Platkiewicz, J. and Brette, R. (2011). Impact of fast sodium channel inactivation on spike threshold dynamics and synaptic integration. PLoS computational biology, 7(5):e1001129.

Popovic, M. A., Foust, A. J., McCormick, D. A., and Zecevic, D. (2011). The spatio-temporal characteristics of action potential initiation in layer 5 pyramidal neurons: a voltage imaging study. The Journal of physiology, 589(17):4167–4187.

Pratt, K. G. and Aizenman, C. D. (2007). Homeostatic regulation of intrinsic excitability and synaptic transmission in a developing visual circuit. Journal of Neuroscience, 27(31):8268–8277.

Pratt, K. G. and Aizenman, C. D. (2009). Multisensory integration in mesencephalic trigeminal neurons in xenopus tadpoles. Journal of neurophysiology, 102(1):399–412.

Pratt, K. G., Dong, W., and Aizenman, C. D. (2008). Development and spike timing–dependent plasticity of recurrent excitation in the xenopus optic tectum. Nature neuroscience, 11(4):467.

Pratt, K. G. and Khakhalin, A. S. (2013). Modeling human neurodevelopmental disorders in the xenopus tadpole: from mechanisms to therapeutic targets. Disease models & mechanisms, pages dmm–012138.

Prinz, A. A., Abbott, L., and Marder, E. (2004a). The dynamic clamp comes of age. Trends in neurosciences, 27(4):218–224.

Prinz, A. A., Bucher, D., and Marder, E. (2004b). Similar network activity from disparate circuit parameters. Nature neuroscience, 7(12):1345.

Reimann, M. W., Nolte, M., Scolamiero, M., Turner, K., Perin, R., Chindemi, G., Dłotko, P., Levi, R., Hess, K., and Markram, H. (2017). Cliques of neurons bound into cavities provide a missing link between structure and function. Frontiers in computational neuroscience, 11:48.

Richards, B. A. and Lillicrap, T. P. (2019). Dendritic solutions to the credit assignment problem. Current opinion in neurobiology, 54:28–36.

Rubinov, M., Sporns, O., Thivierge, J.-P., and Breakspear, M. (2011). Neurobiologically realistic determinants of self-organized criticality in networks of spiking neurons. PLoS computational biology, 7(6):e1002038.

Shen, W., McKeown, C. R., Demas, J. A., and Cline, H. T. (2011). Inhibition to excitation ratio regulates visual system responses and behavior in vivo. Journal of neurophysiology, 106(5):2285–2302.

Spruston, N., Jaffe, D. B., Williams, S. H., and Johnston, D. (1993). Voltage-and space-clamp errors associated with the measurement of electrotonically remote synaptic events. Journal of Neurophysiology, 70(2):781–802.

Stein, B. E., Stanford, T. R., and Rowland, B. A. (2014). Development of multisensory integration from the perspective of the individual neuron. Nature Reviews Neuroscience, 15(8):520.

Stemmler, M. and Koch, C. (1999). How voltage-dependent conductances can adapt to maximize the information encoded by neuronal firing rate. Nature neuroscience, 2(6):521.

Takemura, S.-y. (2014). Connectome of the fly visual circuitry. Microscopy, 64(1):37–44.

Tien, N.-W. and Kerschensteiner, D. (2018). Homeostatic plasticity in neural development. Neural development, 13(1):9.

Titley, H. K., Brunel, N., and Hansel, C. (2017). Toward a neurocentric view of learning. Neuron, 95(1):19–32.

Truszkowski, T. L., Carrillo, O. A., Bleier, J., Ramirez-Vizcarrondo, C. M., Felch, D. L., McQuillan, M., Truszkowski, C. P., Khakhalin, A. S., and Aizenman, C. D. (2017). A cellular mechanism for inverse effectiveness in multisensory integration. Elife, 6.

Turrigiano, G. (2007). Homeostatic signaling: the positive side of negative feedback. Current opinion in neurobiology, 17(3):318–324.

Turrigiano, G. (2011). Too many cooks? intrinsic and synaptic homeostatic mechanisms in cortical circuit refinement. Annual review of neuroscience, 34:89–103.

Venables, W. N. and Ripley, B. D. (2013). Modern applied statistics with S-PLUS. Springer Science & Business Media.

Wu, G.-Y., Malinow, R., and Cline, H. (1996). Maturation of a central glutamatergic synapse. Science, 274(5289):972–976.

Xu, H., Khakhalin, A. S., Nurmikko, A. V., and Aizenman, C. D. (2011). Visual experience-dependent maturation of correlated neuronal activity patterns in a developing visual system. Journal of Neuroscience, 31(22):8025–8036.

Zbili, M., Rama, S., Yger, P., Inglebert, Y., Boumedine-Guignon, N., Fronzaroli-Molinieres, L., Brette, R., Russier, M., and Debanne, D. (2019). Axonal na+ channels detect and transmit levels of input synchrony in local brain circuits. bioRxiv, page 618710.

